# Boosting Transcript Assembly via Delineating Transcript Start and End Sites

**DOI:** 10.1101/2025.10.13.682211

**Authors:** Irtesam Mahmud Khan, Xiaofei Carl Zang, Ange Teng, Tasfia Zahin, Mingfu Shao

**Affiliations:** Department of Computer Science and Engineering, The Pennsylvania State University, 201 Old Main,University Park, 16802, PA, USA; Huck Institutes of the Life Sciences, The Pennsylvania State University, 201 Old Main, University Park, 16802, PA, USA; High Technology High School, 765 Newman Springs Rd, Lincroft, 07738, NJ, USA; Center for Computational and Genomic Medicine, The Children’s Hospital of Philadelphia, 3501 Civic Center Blvd, Philadelphia, 19104, PA, USA

## Abstract

Transcript assembly remains a challenging task despite the development of numerous methods. A major contributor to low assembly accuracy is the difficulty in accurately determining transcript start sites (TSSs) and end sites (TESs), due to the weak and noisy signals typically found in RNA-seq data. We present Telos, a two-stage machine learning framework for precise detection of TSSs and TESs and for transcript ranking. The method takes as input any assembly, typically generated by an existing assembler. In the first stage, Telos scores the TSSs and TESs in the input assembly using a machine learning model trained on a rich set of engineered features. These site-level scores will be passed to the second stage for transcript-level evaluation. In its second stage, Telos scores the entire transcripts by training another model that integrates features of their TSS and TES (including the inferred probabilities from the first stage), along with transcript abundance estimated by the assembler and statistics about exon lengths. We extensively evaluated Telos on ONT (cDNA and direct RNA), PacBio, and Illumina short-read RNA-seq datasets. In all cases, it consistently outperformed baseline methods. Telos is agile, but achieves substantial improvements, demonstrating the value of explicitly modeling TSS and TES, a gap in current transcript assembly tools. Telos can be paired with any assembler to accurately score the assembled transcripts. It is modular, easily extensible to emerging sequencing technologies, and hence we anticipate its broad adoption in transcriptomic studies.

## 1 Introduction

Messenger RNAs (mRNAs) play a central role in celluar functions. They are transcribed from DNA through a complex process, where RNA Polymerase II recognizes proximal promoter regions and synthesizes the precursor mRNA (pre-mRNA) from the transcription start site (TSS) to the transcription end site (TES). Both TSS and TES have important biological significance. Genes with multiple TSS are frequently observed across different organisms [20]. Consequently, this can lead to mRNA isoforms with variable lengths of 5^*′*^ untranslated regions (UTR) in the same gene. Large-scale analysis such as FANTOM CAGE [16] has shown that the majority of mammalian genes contain composite promoters packed with multiple nearby TSSs. This phenomenon can also vary across tissues. Alternative TSSs not only generate different isoforms, but also contribute to the reliability and stability of the mRNA molecule [1]. Alternative TSS usage is commonly observed across diverse cancer types, influencing numerous cancer-related genes, and potentially offering a more accurate prognostic indicator [7]. Similarly to TSS, many genes contain multiple TESs, producing mRNA isoforms with different 3^*′*^ UTR lengths or even alternative last exons. Likewise, this event is recognized as an important regulatory mechanism in gene expression and is implicated in processes from development to cancer. For instance, many genes tend to express shorter 3^*′*^ UTR isoforms in proliferative or cancerous cells, avoiding microRNA sites in longer UTRs [30].

TSS and TES are of highly biological significance and have been extensively studied in various perspectives. In this work, we investigate their link with reference-based transcript assembly, a task referred to as reconstructing the full-length expressed transcripts from RNA-seq alignments. A fully assembled transcript is represented as the genomic coordinates (w.r.t. to the reference genome) of its exons, or equivalently, the TSS, coordinates of splicing junctions, and then the TES. See Figure 1. Transcript assembly methods must therefore identify the TSS, TES, and junctions, and thread them together into complete transcript models. While splicing junctions are relatively easy to detect due to strong signals from reads containing junctions, TSS and TES are much harder to pinpoint due to the lack of such clear indicators. This challenge in accurately identifying TSS and TES is a major contributor to the unsatisfying accuracy of current transcript assembly methods.

**Figure 1.**
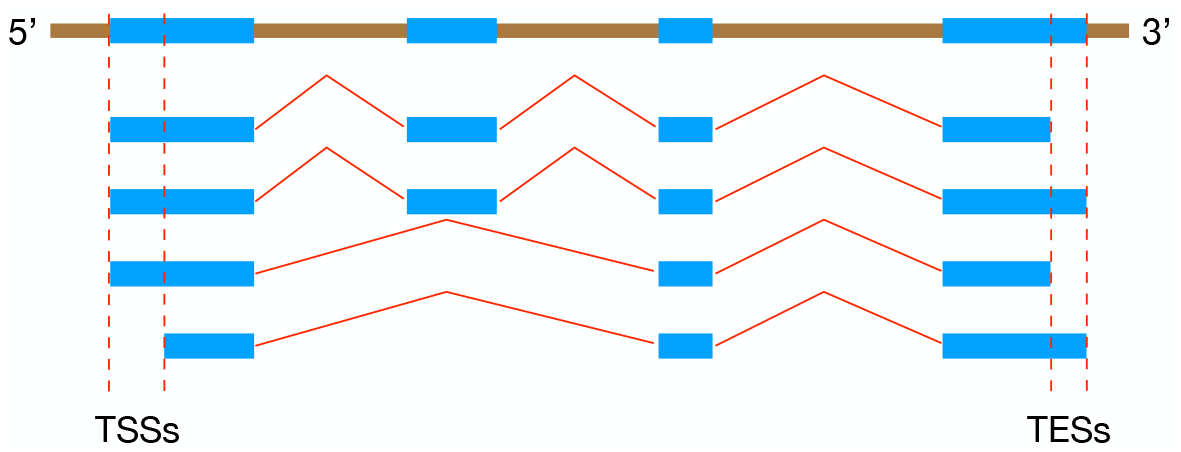
Illustration of alternative TSS and TES. In this example there are two TSSs and two TESs, marked with dashed red lines.

Existing assemblers differ in their approach in determing TSS and TES. State-of-the-art short read-based transcriptome assemblers, such as Scallop2 [31] and StringTie2 [10], used a simple greedy strategy, that only considers TSS and TES as the furthest endpoints of each read cluster. Long-read sequencing offers a compelling advantage—the ability to sequence full-length transcripts from 5^*′*^ cap to 3^*′*^ polyA tail in one read, a significant outperformance in capturing the termini of the molecules compared to short reads [3]. In theory, TSS and TES can be simultaneously delineated for each isoform using full-length long reads. Several computational methods can be used to identify TSS and TES from long-read RNA-Seq reads, but they all, nevertheless, entail compromises [2]. FLAIR relies on annotated TSS positions, trimming or discarding reads that do not align with known start sites; this strategy ensures high accuracy but limits the detection of novel transcript boundaries [27]. TALON clusters read ends to identify novel TSS and TES, filtering out internal priming artifacts at transcript ends [29]. It offers high sensitivity but can produce false positives if thresholds are insufficiently strict. StringTie2 constructs splice graphs to assemble transcripts, refining transcript ends using short-read RNA-seq data [11]. This hybrid approach balances sensitivity and precision effectively but can exclude weakly supported novel ends. Bambu employs a machine-learning model leveraging poly(A) signals and known annotations to validate TSSs/TESs, achieving high precision but potentially omitting genuine low-expression isoforms [4]. IsoQuant identifies TSS and TES through an intron-based transcript graph, selecting the longest read per splice variant to minimize artifacts from truncated reads, resulting in robust sensitivity and precision but occasionally merging biologically distinct isoforms differing only slightly in length [21]. SQANTI3 serves as a post-processing classifier, evaluating transcript completeness based on adapter sequences and poly(A) motifs [17]. Although highly effective for quality control, its accuracy depends strongly on input from other assembly tools. These assemblers employ differing heuristics for identifying TSS and TES, resulting in inconsistent results and reduced overall accuracy (Figure 9). Some of the tools also rely on reference annotation, which is not readily available for novel isoforms and for new organisms.

**Table 1.**
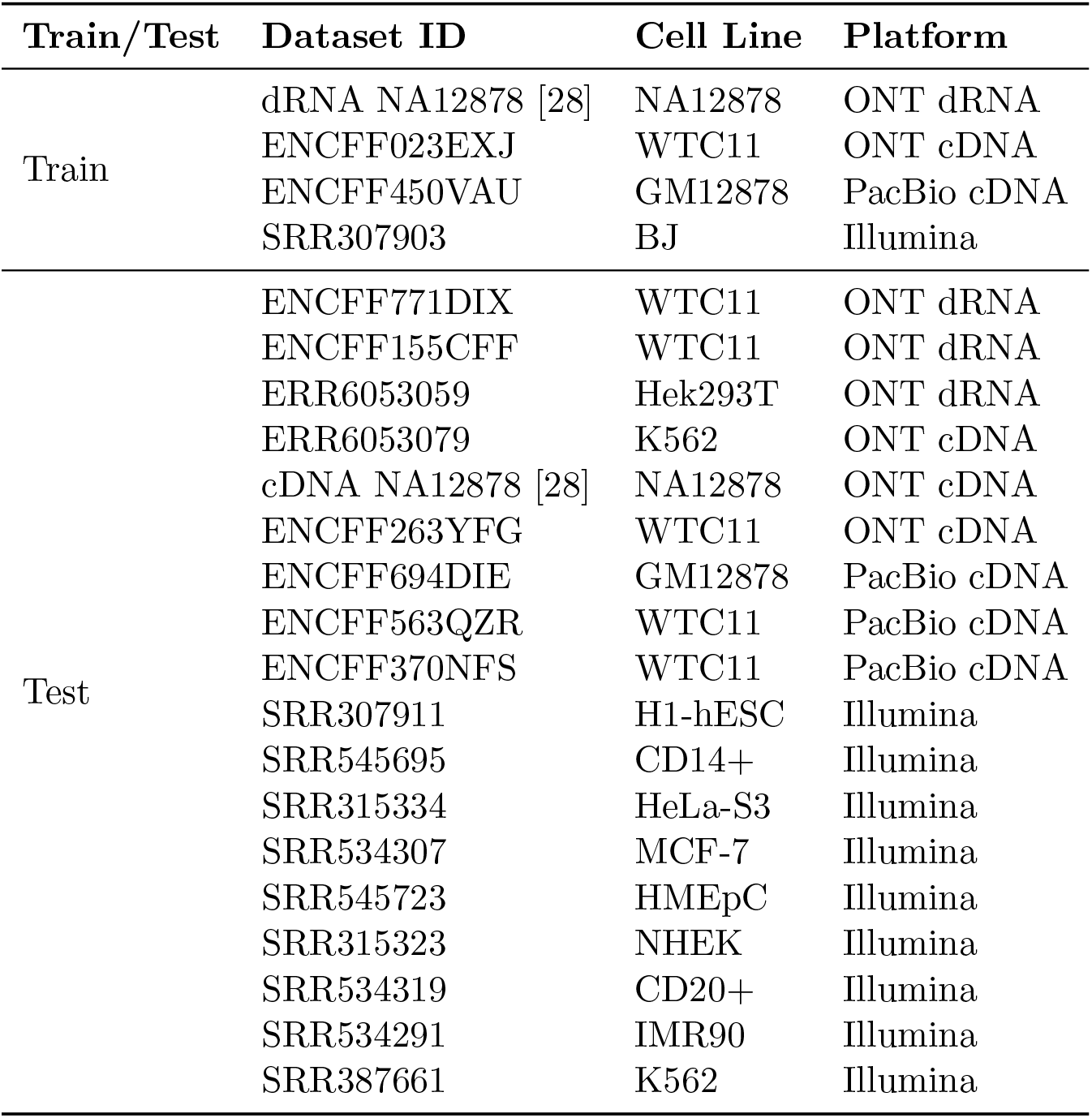
Datasets from four different sequencing technologies were used to train/evaluate our model.

There are specialized sequencing techniques such as CAGE [26], QuantSeq [14], and 3’-end sequencing for polyA sites, that directly detect TSS/TES regions. These methods produce valuable resources including FANTOM [16] and PolyASite databases [15] for research in TSS and TES. Some other studies can predict transcription initiation sites directly using genomic sequences [6, 8]. All sequencing methods measure TSS/TES of the genome in isolation, losing connection to expressed transcripts in a particular sample, and hence not relevant to transcript assembly. DeepBound [23] predicts boundaries of transcripts by utilizing deep convolutional neural fields, but likewise to CAGE and QuantSeq, it neglected connectivity between transcript boundaries and individual isoforms, which obstructs downstream applications.

We present Telos, a machine-learning approach that boosts the assembly accuracy by modeling the TSSs and TESs. Telos takes an assembly (a set of assembled transcripts) from any assembler as input, and predicts a quality score of each individual transcript and its TSS and TES. Telos uses a rich set of features we engineered to comprehensively describe the TSS and TES of the transcripts. A model is then trained using these features to infer the quality score. It is important to note that while Telos is trained using genome annotations as ground truth, it does not rely on annotations or genomic coordinates during prediction. This enables Telos to generalize beyond known annotations, effectively identifying novel TSSs/TESs and facilitating its use across diverse or newly sequenced organisms. Through extensive experiments, we demonstrate that Telos consistently outperforms baseline approaches across a variety of sequencing datasets.

## 2 Results

### 2.1 Overview of Telos

Telos is implemented as a two-stage classification framework. See Figure 2. In the first stage, Telos takes as input an aligned BAM file containing sequencing reads and a GTF file containing assembled transcripts. From the GTF file, Telos extracts candidate transcription start sites (TSSs) and transcription end sites (TESs), and computes a set of engineered features for each site based on the distribution of nearby reads. These site-level features are then evaluated using a machine learning classifier (Random Forest or XGBoost) to estimate the probability that each TSS or TES is correct. In the second stage, Telos aggregates site-level information to evaluate transcripts as a whole. For each transcript, the features generated for its start and end sites, together with their predicted correctness probabilities, are concatenated. Additional transcript-level features are also incorporated, including basic statistics such as transcript length, minimum/maximum/mean exon length, and the lengths of the first and last exons. The resulting feature vector is then provided to an XGBoost classwifier, which assigns a probability score reflecting the overall correctness of the predicted transcript.

**Figure 2.**
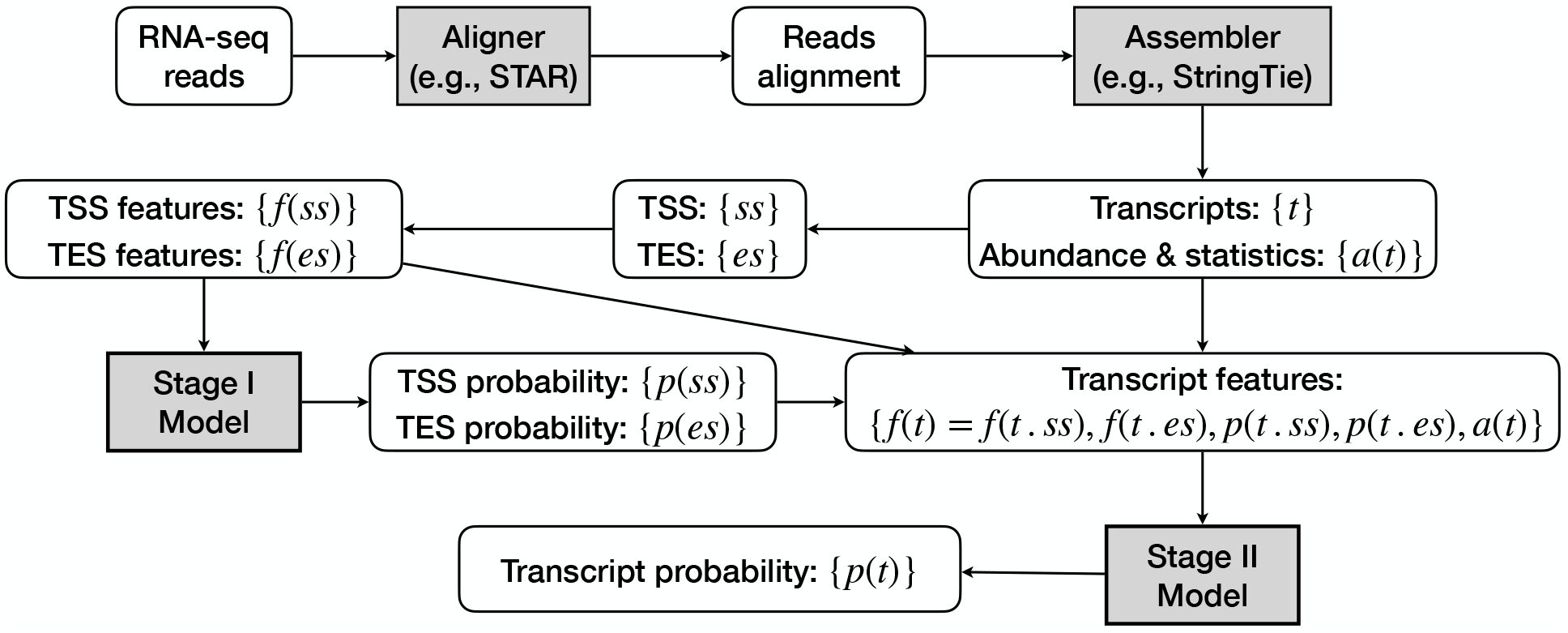
Overall pipeline of Telos.

### 2.2 Dataset

To train and evaluate Telos, we compiled a diverse set of transcriptomic datasets spanning multiple sequencing technologies and cell types. Four datasets were used for training: cDNA (ENCFF023EXJ) datasets from the WTC11 cell line, direct RNA (dRNA) from NA12878 cell line, a PacBio human GM12878 cell line data (ENCFF450VAU) from the ENCODE project, and an Illumina short-read RNA-seq dataset (SRR307903). The dRNA data provides native RNA molecules with preserved 3’ ends, while the corresponding cDNA data offers higher throughput but is more susceptible to 5’ truncation artifacts. The PacBio dataset represents full-length transcript sequencing with high accuracy, and the short-read RNA-seq dataset was included to ensure that the training procedure generalizes across multiple sequencing modalities.

To test the performance of Telos, we evaluated it on eighteen independent datasets that were not used during training. These include nine long read datasets and another nine short read datasets.

These test datasets encompasses diverse cell types and sequencing batches, allowing us to assess the robustness of our method under varied conditions.

### 2.3 Scoring TSS and TES Sites

Although the primary goal of this study is to improve transcript assembly by scoring and ranking complete transcript models, the first stage of Telos focuses on independently evaluating TSS and TES sites produced by different assemblers. In this stage, we assessed the performance of classifier-assembler combinations using precision-recall (PR) curves, as shown in Figures 3 to 6. For comparison, baseline PR curves were generated by ranking candidate sites solely on transcript abundance, using the mean abundance of the assembled transcripts that contributed each site. All reported results are based on the validation sets of the four training datasets. To avoid data leakage, all models presented in this study were trained exclusively on chromosomes 1–10 and validated on the remaining chromosomes.

**Figure 3.**
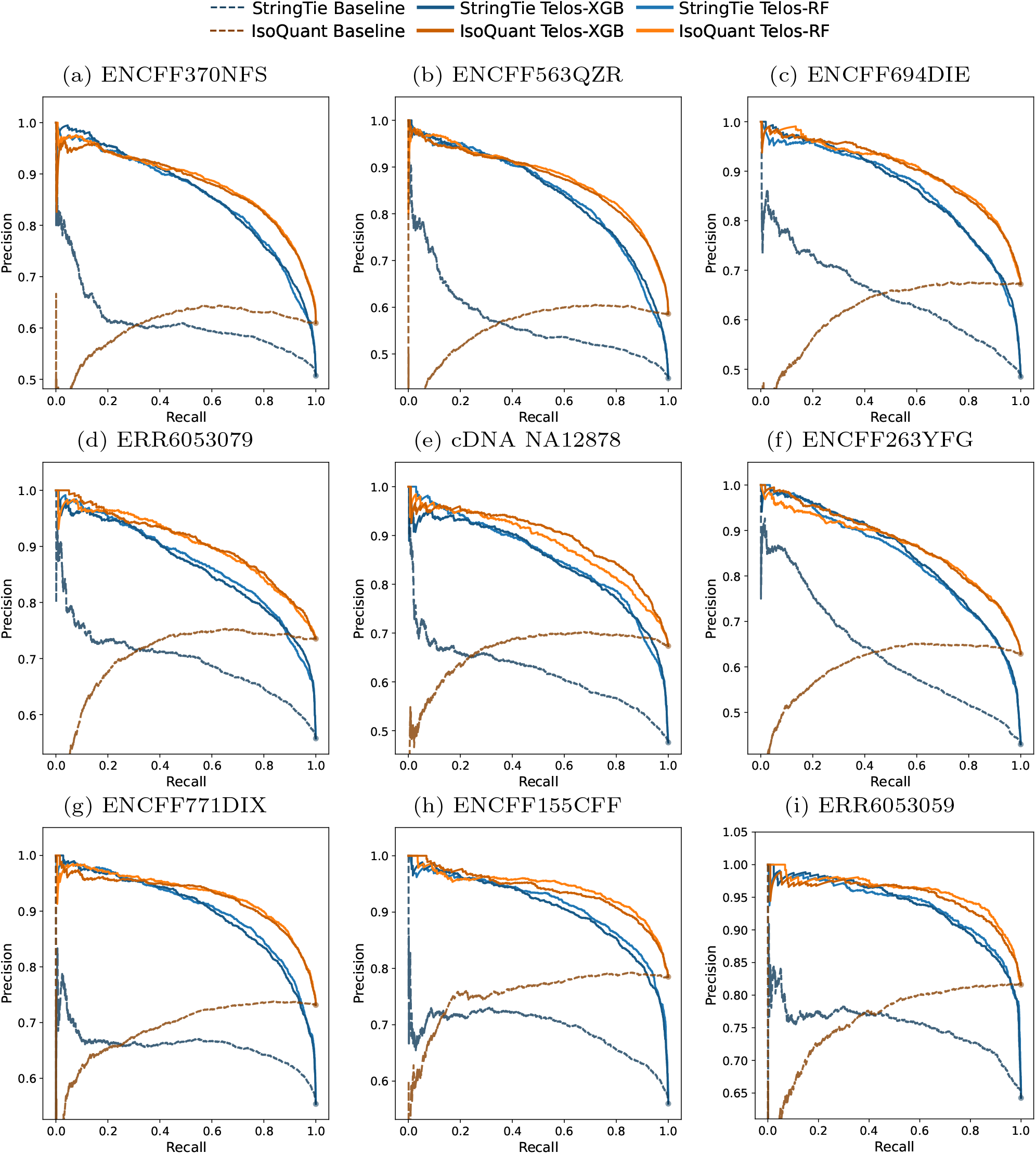
Precision-recall (PR) curves for TSS classification across multiple long read datasets. Figure (a)-(c) are results for pacbio datasets, (d)-(f) are from cDNA datasets and (g)-(i) are from dRNA dataset. Each panel shows PR curves for TSS prediction for a particular test dataset. Different shades of Blue (StringTie) and Orange (IsoQuant) indicates the underlying assembler used. Each dashed line corresponds to an assembler’s baseline predictions, whereas the solid line indicates Telos’s predictions.

**Figure 4.**
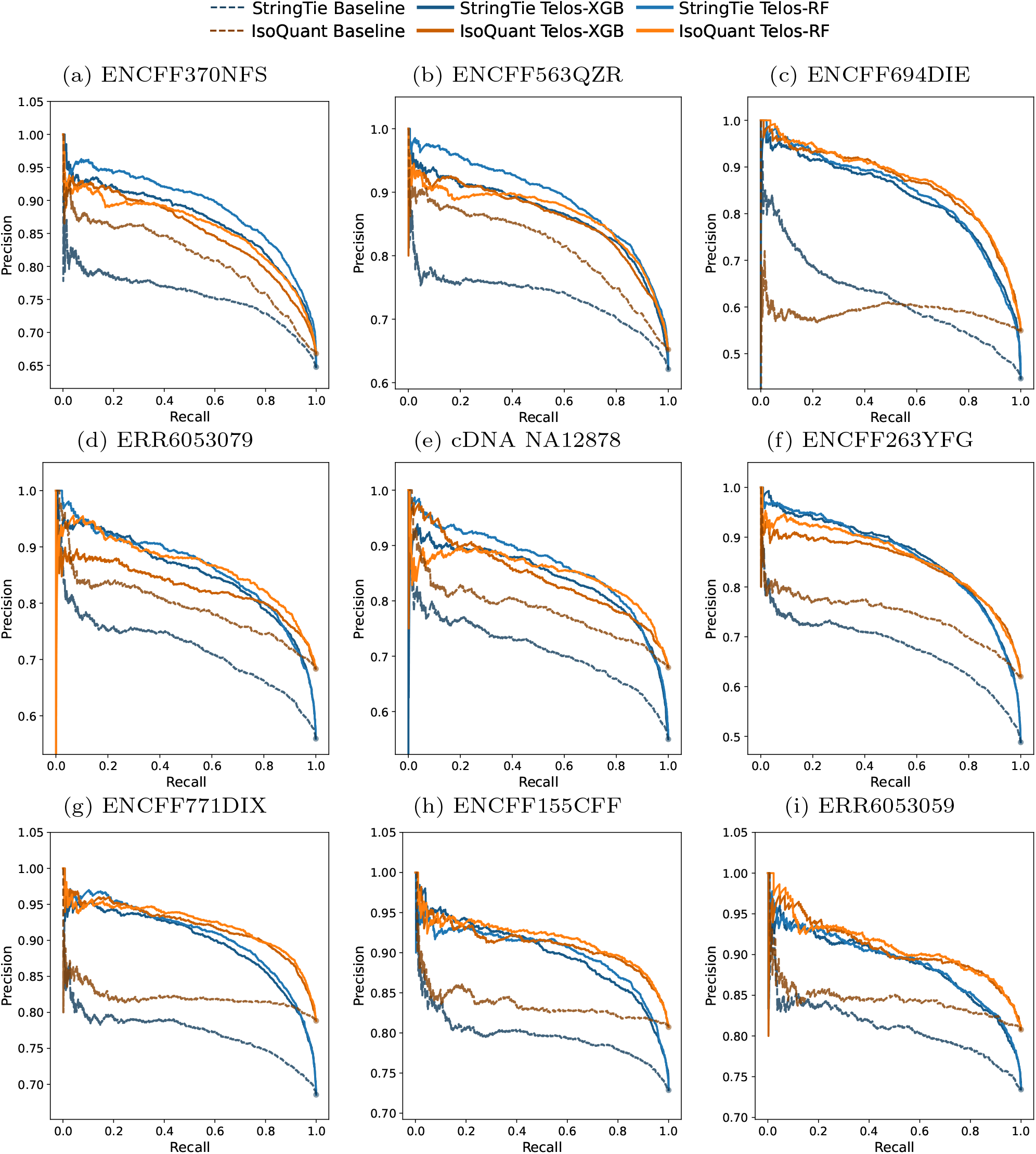
Precison-recall (PR) curves for TES classification across multiple long read datasets. Figure (a)-(c) are results for pacbio datasets, (d)-(f) are from cDNA datasets and (g)-(i) are from dRNA dataset. Each panel shows PR curves for TES prediction for a particular test dataset. Different shades of Blue (StringTie) and Orange (IsoQuant) indicates the underlying assembler used. Each dashed line corresponds to an assembler’s baseline predictions, whereas the solid line indicates Telos’s predictions.

**Figure 5.**
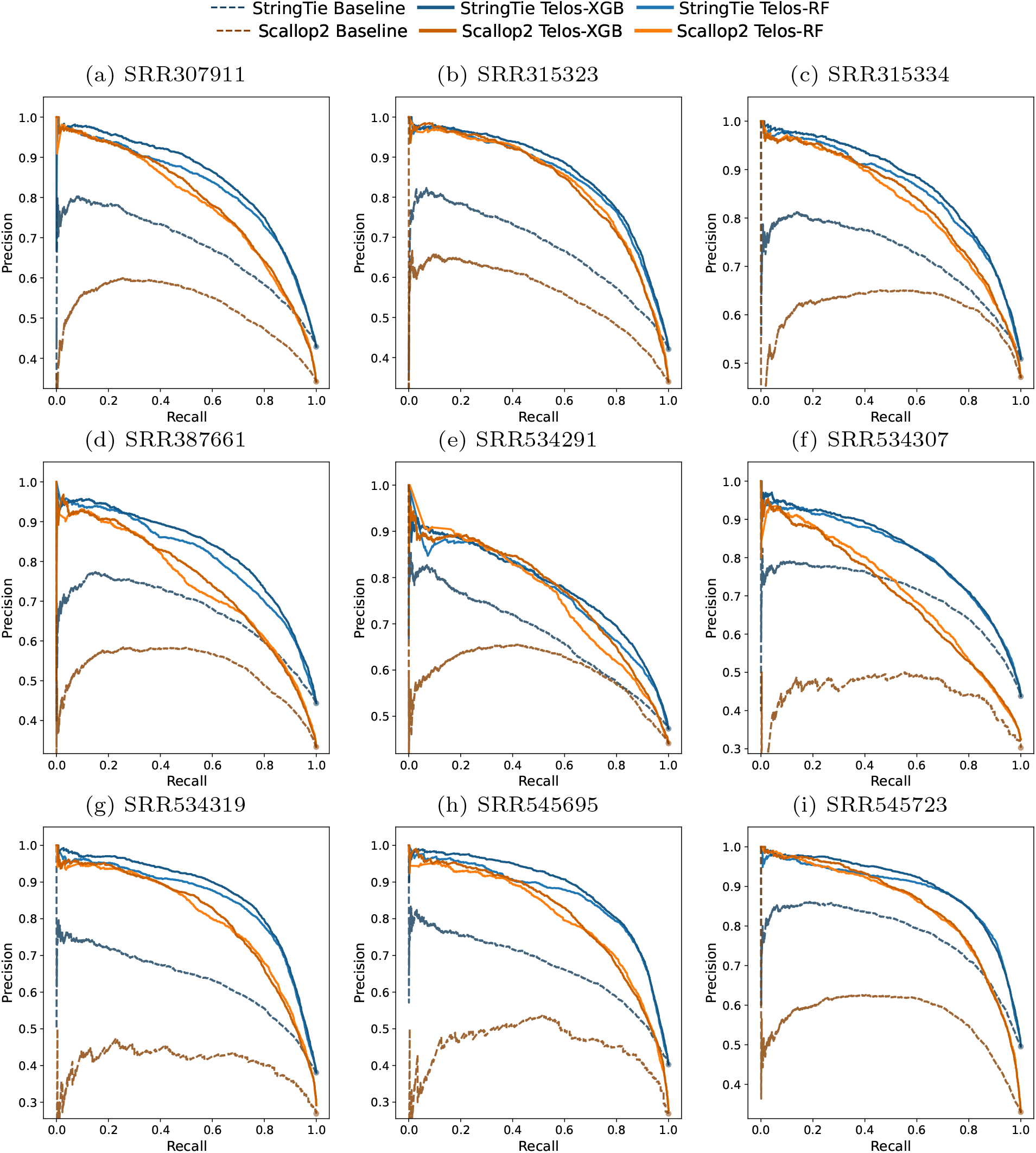
Precison-recall (PR) curves for TSS classification across multiple short read datasets. Each panel shows PR curves for TSS prediction for a particular test dataset. Different shades of Blue (StringTie) and Orange (Scallop2) indicates the underlying assembler used. Each dashed line corresponds to an assembler’s baseline predictions, whereas the solid line indicates Telos’s predictions.

**Figure 6.**
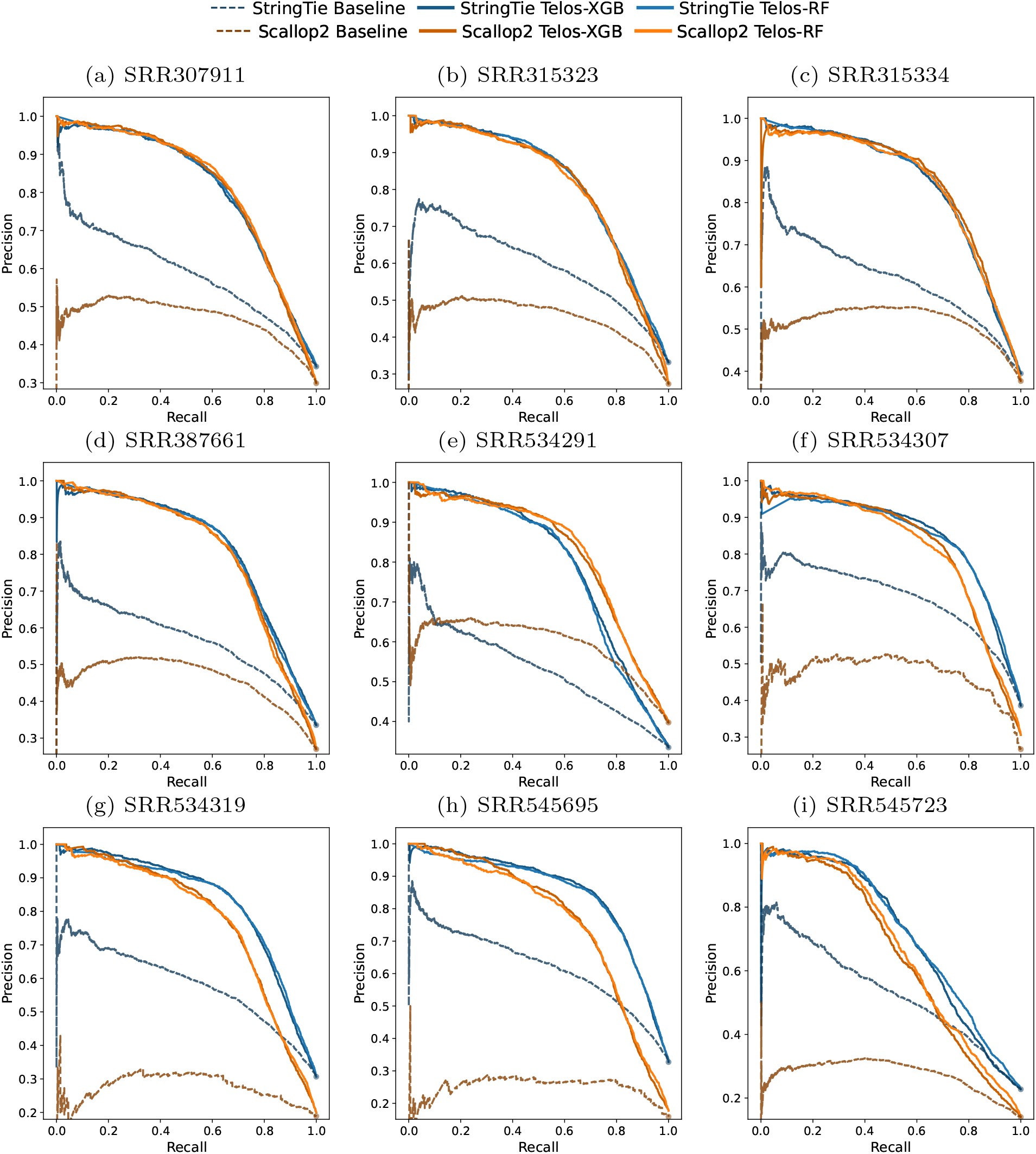
Precison-recall (PR) curves for TES classification across multiple short read datasets. Each panel shows PR curves for TES prediction for a particular test dataset. Different shades of Blue (StringTie) and Orange (Scallop2) indicates the underlying assembler used. Each dashed line corresponds to an assembler’s baseline predictions, whereas the solid line indicates Telos’s predictions.

To compare the area under precision-recall curve (AuPR) values for each test dataset, we plotted them as barplots in Figures 7 and 8. Telos demonstrates substantial improvements over baseline coverage-based methods across all dataset types and assemblers. For TSS detection, Telos achieves average improvements of 25.8-50.1% across dataset groups, with the highest gains observed for PacBio datasets (40.4% with StringTie, 50.1% with IsoQuant) and short-reads datasets using Scallop2 assemblers (60.0%). For TES detection, improvements range from 10.0-108.0%, with short-reads datasets showing the most dramatic enhancements (38.3% with StringTie, 108.0% with Scallop2). Notably, dRNA datasets exhibit more modest improvements (10.0-31.1%), while cDNA and PacBio datasets show consistent gains of 19.9-40.4%. These results demonstrate that Telos provides robust performance improvements across diverse sequencing technologies and assembly approaches, with particularly strong benefits for short-read sequencing data.

**Figure 7.**
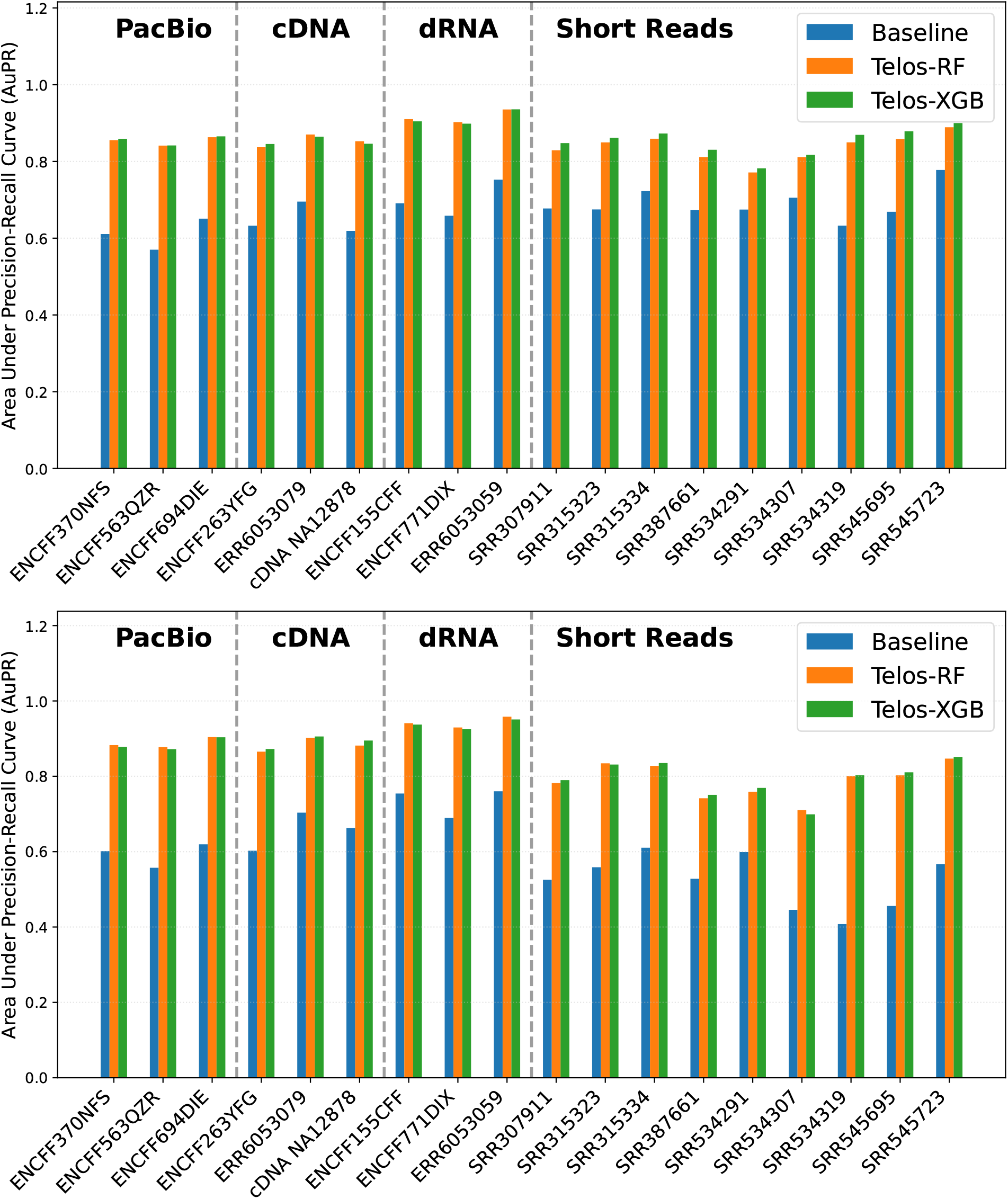
Comparison of baseline assembler and Telos models on TSS prediction performance. Each bar represents the area under the precision-recall curve (AuPR) for a single dataset, evaluating transcription start site (TSS) prediction accuracy. Blue bars correspond to the baseline assembler, orange bars to the Telos–Random Forest model, and green bars to the Telos–XGBoost model. Bars are grouped according to the underlying sequencing technology (PacBio, ONT direct RNA, ONT cDNA, and short-read). The top panel uses StringTie as the baseline assembler, while the bottom panel shows results for IsoQuant and Scallop2.

**Figure 8.**
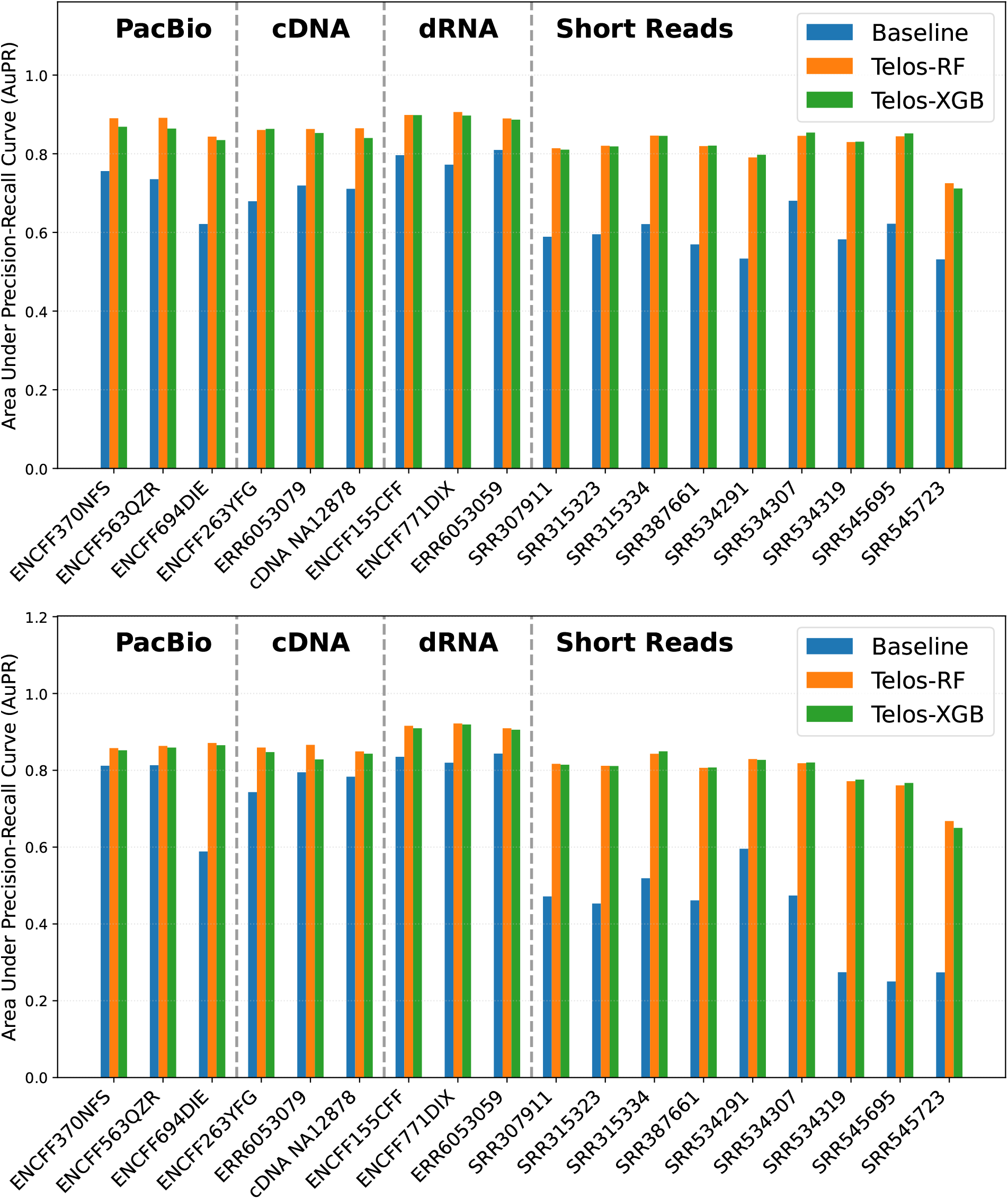
Comparison of baseline assembler and Telos models on TES prediction performance. Each bar shows the area under the precision-recall curve (AuPR) for transcription end site (TES) prediction across different datasets. As in Figure 7, blue bars denote the baseline assembler, orange bars the Telos–Random Forest model, and green bars the Telos–XGBoost model. Bars are grouped by sequencing technology (PacBio, ONT direct RNA, ONT cDNA, and short-read). The top and bottom panels correspond to results using StringTie and other assemblers (IsoQuant, Scallop2), respectively.

Telos consistently achieves a significantly larger area under the precision-recall (PR) curves across all datasets, demonstrating clear improvements over the baseline methods. Across all long-reads datasets, transcription start site (TSS) and transcription end site (TES) candidates generated by IsoQuant consistently outperformed those from StringTie. A major contributing factor is that the IsoQuant baseline itself performs better than StringTie, particularly in the high-recall region. This trend suggests that IsoQuant makes more precise predictions overall, whereas StringTie generates a more diverse but recall-rich set of candidates. For short-reads data, StringTie consistently outperforms Scallop2 in terms of TSS/TES classification accuracy. Interestingly, we observe an atypical behavior in the IsoQuant baseline predictions for TSS classification: as recall decreases with transcript abundance, precision also decreases, rather than increasing as would normally be expected. A similar pattern is observed for short-reads datasets, although the effect is less pronounced. In contrast, for StringTie predictions, precision does increase as recall decreases; however, the gains are often negligible in the high-recall region. Taken together, these results indicate that even for highly expressed transcripts, assemblers frequently fail to accurately predict TSS (and, to a lesser extent, TES) positions. Telos, by contrast, achieves consistently superior performance compared to all baselines, with the most pronounced improvements in the high-recall region across all datasets.

Importantly, both Random Forest and XGBoost classifiers produce nearly identical results across datasets and assemblers, suggesting that classifier choice has minimal impact when supplied with a well-engineered feature set.

The TSSs/TESs predicted by different assemblers vary substantially, highlighting the low accuracy of existing methods. In Figure 9 we compare the Jaccard similarity of TSSs/TESs predicted by the two assemblers before and after applying Telos to filter false positives. Across all datasets, concordance between the two assemblers increases markedly after filtering. This provides strong evidence that the majority of inconsistent predictions are attributable to spurious endpoints, which Telos effectively identifies and removes.

Collectively, these findings demonstrate that the first-stage Telos model reliably discriminates between true and false TSSs/TESs across platforms and assemblers, thereby establishing a solid foundation for more accurate transcript-level scoring in the second stage.

### 2.4 Scoring Transcripts

We now present our results that Telos improves transcript assembly. It is well known that the predicted abundance (by an assembler) highly relates to their correctness [22, 31, 10, 19, 13, 21]. Hence, ranking transcripts by their abundances are a common practice in transcriptomic research. The precision-recall curves, by varying the abundance, is often drawn to compare different assemblers. This serves as our baseline method. Recall that Telos eventually assigns a confidence score to each transcript in the input assembly. We draw the precision-recall curve by varying this score to compare with the baseline methods. We also report the area under the PR curves for quantitative evaluations. In all methods, ground-truth are again defined using the annotation, where we labeled transcripts as true if they matched a known isoform in the annotation and false otherwise. Because the entire annotation was treated as the “true set”, while the transcripts expressed in a given RNA-seq sample typically represent only a small subset of the annotation, recall values are generally low across all methods; nevertheless, the relative recall among methods still provides a meaningful and fair comparison of their performance.

We compare the precision-recall curves in Figures 10 and 11. Telos consistently outperformedthe baseline models across nearly all datasets. The only exception was observed for the dRNA IsoQuant dataset, where Telos achieved performance comparable to the baseline. Interestingly, for this same dataset, Telos exhibited markedly stronger results in Stage I, indicating a potential inconsistency that we further discussed in Section 3.

**Figure 9.**
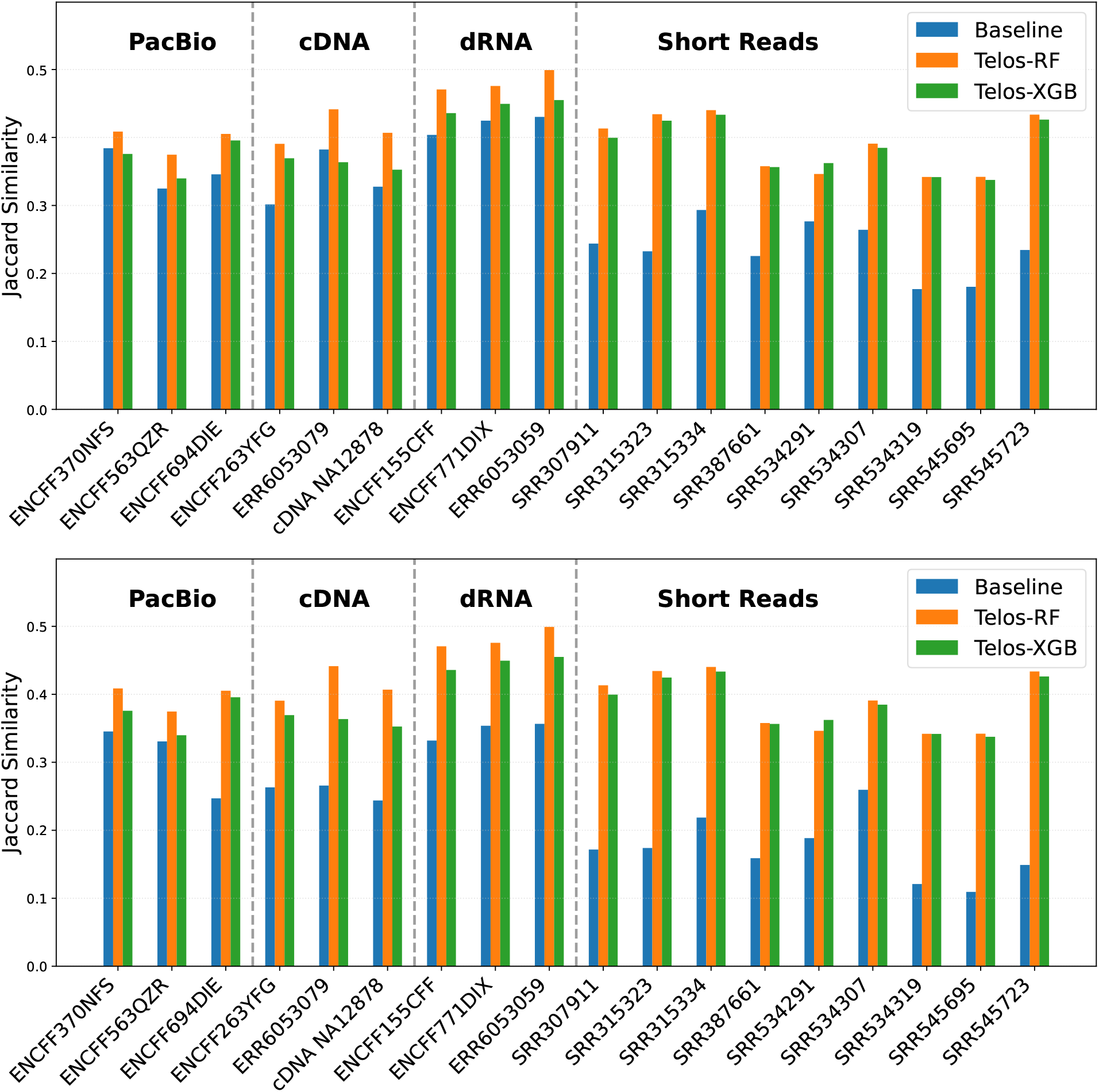
Jaccard similarity of predicted TSS and TES between the two assemblers. The top and bottom panel shows results for TSS and TES respectively. The baseline similarities are shown in blue. The two additional bars indicate the results after removing false positive sites based on Stage I model predictions from Telos. Each group of bars represent a particular sequencing technology as indicated by the label. For the purpose of this analysis, two predicted sites were considered concordant if their genomic coordinates fell within ±50 bp of each other.

**Figure 10.**
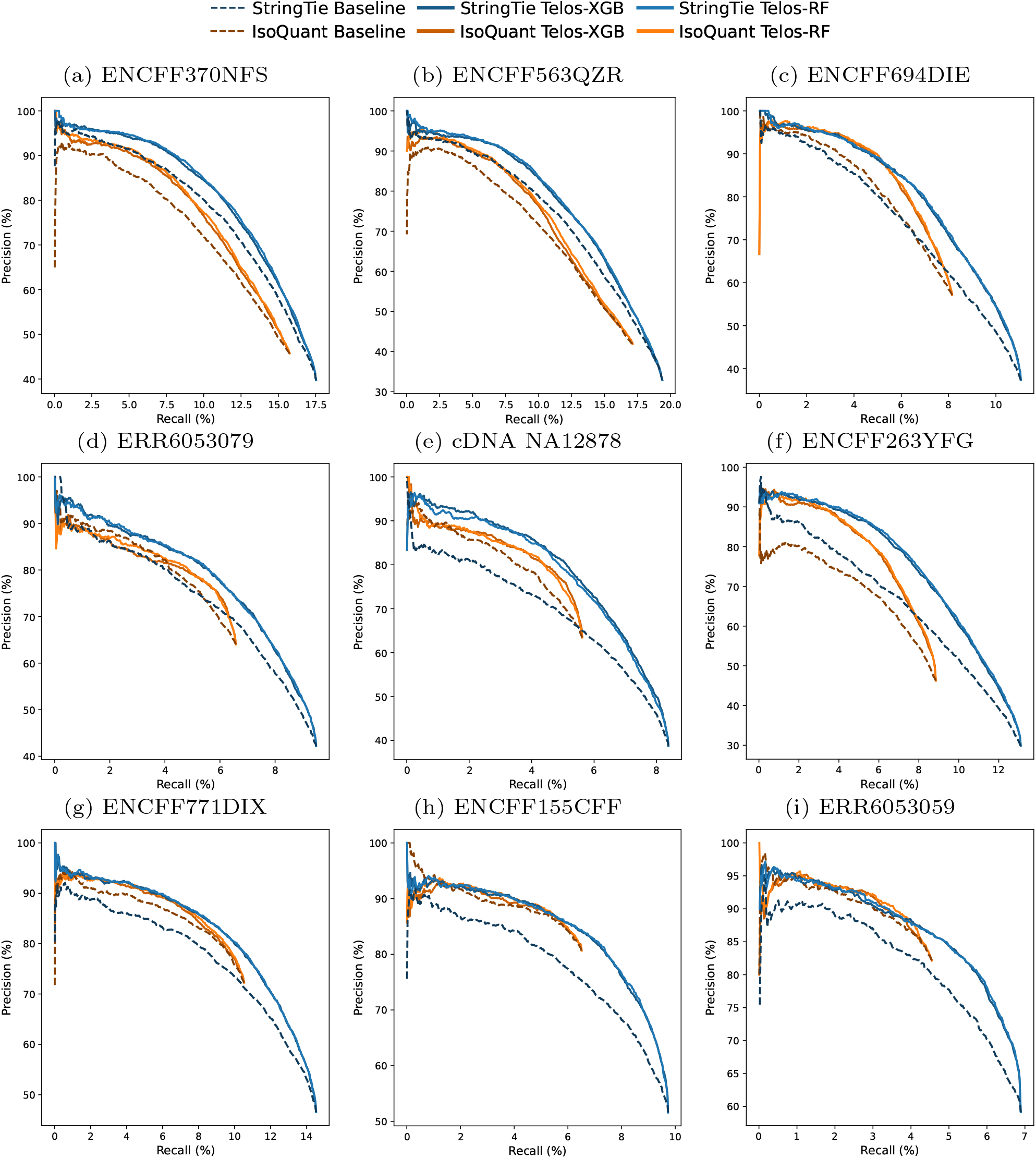
Comparison of precision-recall (PR) curves for transcript scoring on long-reads data. Each row corresponds to a different sequencing technology, Figure (a)-(c) are results for pacbio datasets, (d)-(f) are from cDNA datasets and (g)-(i) are from dRNA dataset. Results for StringTie and IsoQuant are shown in Blue and Orange, respectively. Dashed lines represent the baseline PR curves for the assemblers. Solid lines correspond to model predictions from the Stage I classifiers, with darker and lighter shades denoting XGBoost and Random Forest, respectively. The Stage II model is consistently configured to use LightGBM.

**Figure 11.**
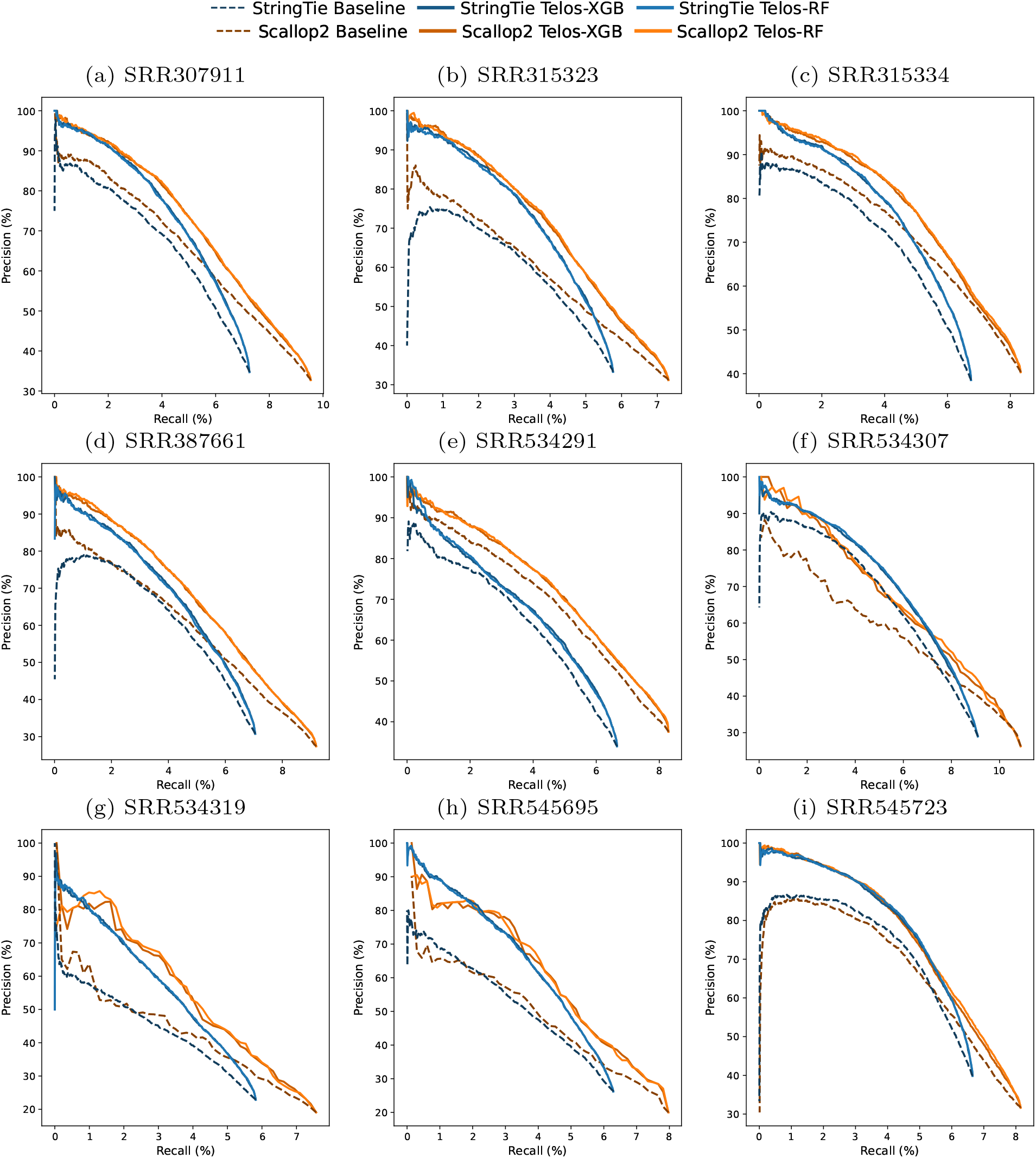
Comparison of precision-recall (PR) curves for transcript scoring on short-reads data. Results for StringTie and Scallop2 are shown in Blue and Orange, respectively. Dashed lines represent the baseline PR curves for the assemblers. Solid lines correspond to model predictions from the Stage I classifiers, with darker and lighter shades denoting XGBoost and Random Forest, respectively. The Stage II model is consistently configured to use LightGBM.

We report the corresponding area under the precision-recall curves in Figure 12. The results demonstrate that Telos consistently enhances transcript assembly across all test datasets. Given the extensive number of transcripts in the human genome, even a marginal improvement in precision can translate to the removal of hundreds of false positives.

**Figure 12.**
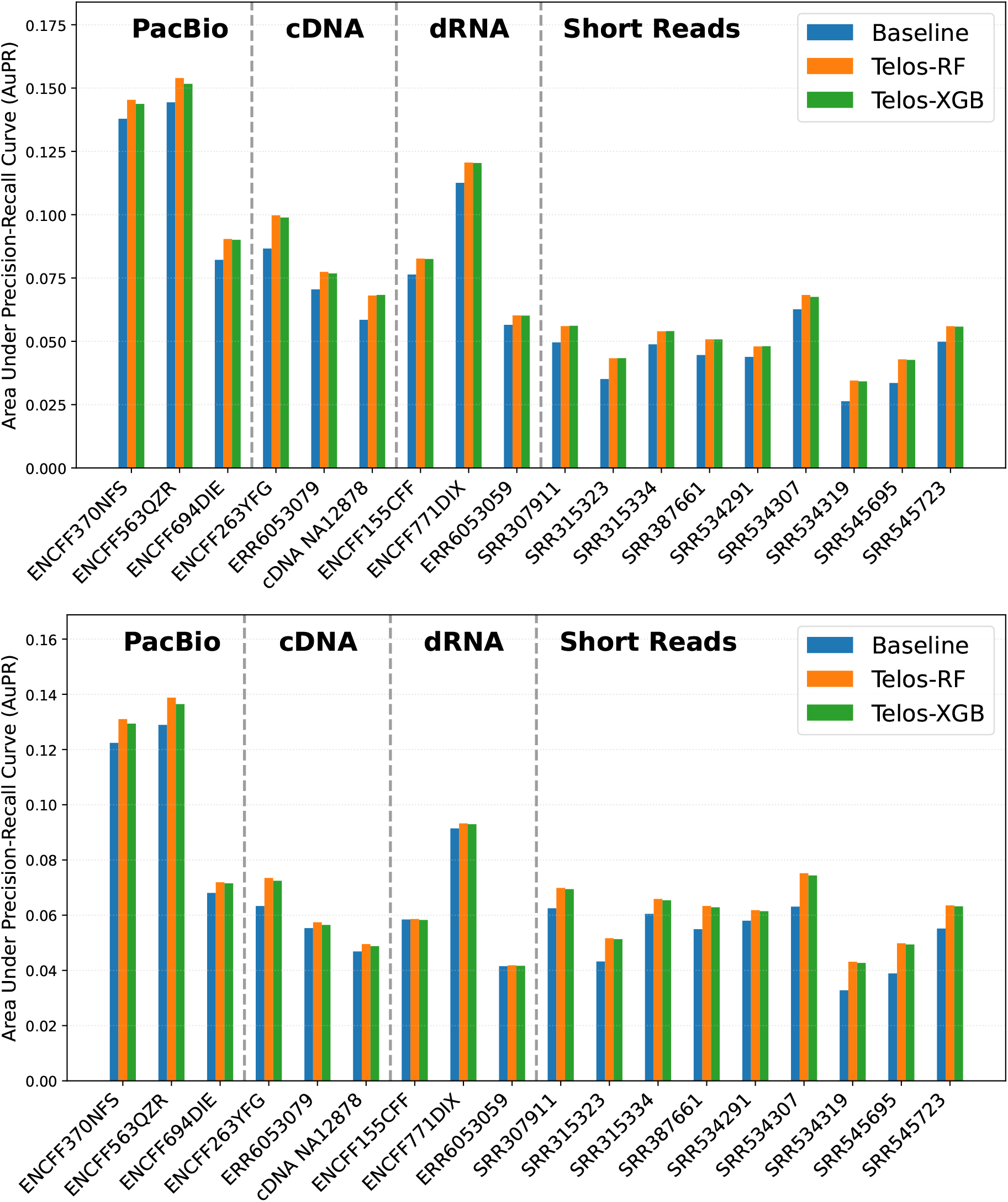
Comparison between the baseline predictions and Telos in terms of area under precision-recall curve (AuPR) on predicting transcripts is presented in this barplot. The top and the bottom panel presents results associated with StringTie and IsoQuant (for long reads) and Scallop2 (for short reads) baseline assemblies respectively. The Blue bars represent the baseline, whereas the Orange and Green bars represent the AuPR values of the Telos predictions with Stage I Random Forest and XGBoost model respectively. The AuPR values were calculated considering the reference annotations as the ground truth. Each group of bars represent a particular sequencing technology as indicated by the label. Telos performs better than the baseline across all test datasets.

The performance gains were most prominent in long-read cDNA datasets and short reads datasts, where inaccurate TSSs/TESs and misassembled isoforms are common. In particular, our method showed improved early precision, meaning it more effectively ranked true transcripts higher than competing approaches. This behavior is especially valuable in downstream workflows such as isoform quantification, where filtering high-confidence transcripts is often essential. The integration of site-derived features from Stage I enabled the model to distinguish well-formed transcript structures even in the absence of full-length support.

The improvement over IsoQuant assemblies is smaller than that over StringTie assemblies on long-reads datasets. One contributing factor is that IsoQuant inherently predicts fewer transcripts and tends to be conservative in its reconstructions. As a result, its output is already more precise, leaving less room for scoring gains. Also, IsoQuant is optimized for long-read input and often excludes low-confidence transcript fragments that StringTie may retain, leading to a naturally lower false positive rate prior to classification. StringTie, on the other hand, typically exhibits higher recall but lower precision, especially in complex loci or low-coverage regions. This makes it more susceptible to spurious transcript models that can be effectively filtered by Telos. The higher gain observed in StringTie-derived assemblies reflects Telos’ ability to distinguish high-confidence models from noise when the initial output is less conservative.

## 3 Discussion

Telos proves that explicitly modeling TSS/TES is critical for improving transcript assembly. Despite using a straightforward approach—classic models trained on engineered tabular features—Telos consistently outperforms across diverse datasets. This success highlights the untapped potential of this direction and opens the door for further advancements in transcript assembly through refined modeling of transcript start and end sites.

Telos can be improved in several ways. First, Telos currently relies on curated annotations (e.g., RefSeq, GENCODE, Ensembl) to label training data, which may omit biologically valid but unannotated. Future work could address this by incorporating semi-supervised learning techniques or reference-free validation strategies, such as clustering signal consistency across replicates or using orthogonal experimental evidence like CAGE or PolyA-seq.

Second, while Telos supports all major long-read sequencing protocols, direct RNA (dRNA) sequencing remains a particular challenge. Our analyses show that Telos achieves the smallest performance gains on dRNA datasets for transcript-level classification, although it continues to outperform baseline models in predicting TSS/TES. This discrepancy likely reflects the nature of dRNA sequencing: because the technology captures full-length RNA molecules directly, assemblers are naturally more effective at identifying true transcript start and end sites, leaving Telos with limited scope for further improvement at this stage. By contrast, the low throughput and noisier error profile of dRNA data often lead to inaccurate exon-intron structures, which Telos cannot fully resolve in Stage II, even when endpoints are refined correctly.

Several additional factors may contribute to the reduced transcript-level performance of Telos on dRNA data. First, dRNA reads are more error-prone than cDNA-based protocols, with higher indel rates and homopolymer-associated errors, which may obscure key alignment-derived features used in Stage I. Second, dRNA coverage is often uneven across transcripts, especially at the 5’ ends, complicating both TSS detection and downstream isoform scoring. Third, biases specific to dRNA (e.g., read truncation, modified bases affecting basecalling and alignment) may limit the discriminative power of the exon-length statistics and endpoint-derived features used in Stage II.

Together, these observations suggest that while Telos reliably identifies true TSS/TES in dRNA datasets, transcript-level scoring requires features that better capture isoform complexity under the constraints of low throughput and high error rates. Addressing these limitations—for example, by incorporating dRNA-specific error models, leveraging modification-aware alignments, or integrating orthogonal evidence such as short-reads data—may enhance the effectiveness of Telos on dRNA data.

Another challenge stems from the dependence on transcriptome assemblers to define candidate TSS and TES positions. As shown in Figure 9, StringTie and IsoQuant often yield divergent candidate sets, with substantial disagreement and non-overlapping sites. These results suggest that further work is needed to improve the upstream candidate generation step, potentially by integrating evidence across multiple tools or incorporating read-level signals directly.

Previous studies have shown that soft-clipping patterns are found in transcript start/end sites in ONT reads [25]. Based on this observation, our tool incorporates multiple features related to soft clip. Our analysis of feature importance also validates this phenomenon, as mutliple soft clip derived features are consistently considered one of the top-ranked features by the Random Forest model. A study by Workmen et al. [28] shows that sharp increases in coverage and read start densities are typically observed at TSSs, while coverage drop-offs and poly-A read ends cluster at TESs; hence, it correlates with our feature ranking where coverage and read density are always one of the top features. We will take into these prior knowledges into account and model the corresponding distributions in the next development of Telos.

## 4 Conclusion

We present Telos, a machine learning-based framework that scores TSS/TES sites and entire transcripts. Our results show that Telos consistently improves transcript assembly across ONT cDNA, ONT dRNA, PacBio Iso-Seq, and Illumina short-read RNA-seq datasets. Telos is modular, compatible with standard RNA-seq alignment and assembly pipelines. It integrates easily with existing tools like StringTie, IsoQuant, and Scallop2, offering a principled post-scoring step for site refinement and transcript ranking. Given its extensibility, Telos can be adapted or re-trained to emerging sequencing technologies and used in downstream tasks such as isoform filtering, expression analysis, and transcript model validation.

## 5 Methods

Telos is a two-stage machine learning method designed to improve the identification of transcript start sites (TSSs) and transcript end sites (TESs) in transcript assemblies. Telos infers TSS and TES locations directly from features derived from aligned RNA-seq reads. The framework is modular and general-purpose, compatible with transcripts assembled from a variety of sequencing platforms. The core idea behind Telos is to treat TSS and TES detection as a supervised classification problem, decoupled from the transcript assembly process itself. Given any assembled assembly and its corresponding aligned reads, Telos extracts sequence- and alignment-based features around TSSs/TESs and uses these to train classifiers that score each TSSs/TESs as likely true or false. These site-level predictions constitute the Stage I of the framework. In the Stage II, Telos aggregates the TSS and TES prediction scores and additional transcript-level features to estimate the overall quality of an assembled transcript. This transcript-ranking model is trained to distinguish between biologically accurate transcript models and likely artifacts, using only features derived from the input assembly and read alignments. The output is a ranked list of assembled transcripts with confidence scores that can be used independently or as input to downstream analyses such as differential isoform usage, gene model refinement, or expression filtering.

### 5.1 Transcript Assembly from RNA-seq Alignments

Telos is designed to operate on data constructed from a wide range of RNA-seq technologies, including Oxford Nanopore Technologies (ONT) cDNA and direct RNA sequencing, Pacific Biosciences (PacBio) Iso-Seq, and Illumina short-read sequencing. Regardless of platform, the pipeline requires two primary inputs: (1) a set of aligned RNA-seq reads in BAM format, and (2) an assembly in GTF format. For long-reads data, input reads are aligned to a reference genome using Minimap2 [12] in spliced alignment mode (-ax splice) with recommended settings for the specific platform. Alignments are strand-aware and preserve soft-clipped bases to enable accurate characterization of TSSs/TESs. Transcripts are assembled using StringTie (version: 3.0.0) and IsoQuant (version: 3.6.3), two representative tools optimized for long-read reconstruction. Both assemblers generate full-length transcript models based on spliced alignments. For short-reads data, we use the HISAT2 [9] aligner to map reads to the genome and assemble transcripts using both StringTie and Scallop2 (version: 1.1.2), two widely-used short-read transcriptome assemblers.

For both short and long reads, Telos processes each transcript from the input GTF file to extract its 5^*′*^ and 3^*′*^ ends (based on strand orientation), forming the initial pool of candidate TSS and TES. Transcripts lacking strand assignment are excluded from analysis. To derive candidate TSSs/TESs from the assembled GTF files, we utilized the GTFFormat tool from the RNASeq-Tools [24] suite, which parses GTF records to identify transcript start and end coordinates. The resulting candidate sites are processed independently for TSS and TES classification in the subsequent stages of the pipeline.

### 5.2 Stage I: TSS and TES Scoring

In the first stage of the Telos, we aim to assess the correctness of TSSs and TESs as predicted by a given assembly. Two independent models (supervised machine learning models) are trained to classify candidate TSSs and TESs, respectively, based on a rich set of alignment-derived and sequence-context features.

#### Feature Extraction from Aligned Reads

For each candidate site, Telos computes a set of engineered features from the aligned reads in the corresponding BAM file. These features are designed to capture local patterns of read density, coverage shifts, splicing context, soft clips and alignment quality, all of which are informative for TSS/TES classification. Feature extraction is performed within a fixed genomic window centered on each candidate site (by default *±*100 nt). These features are computed using a custom Python module built on pySAM [5], and are fully assembler- and platform-agnostic. Besides, no genomic coordinates or annotation-specific features were provided to Telos for training. See Figure 13 for an example in which we illustrate three features. An overview of all the features are described in Appendix A.

**Figure 13.**
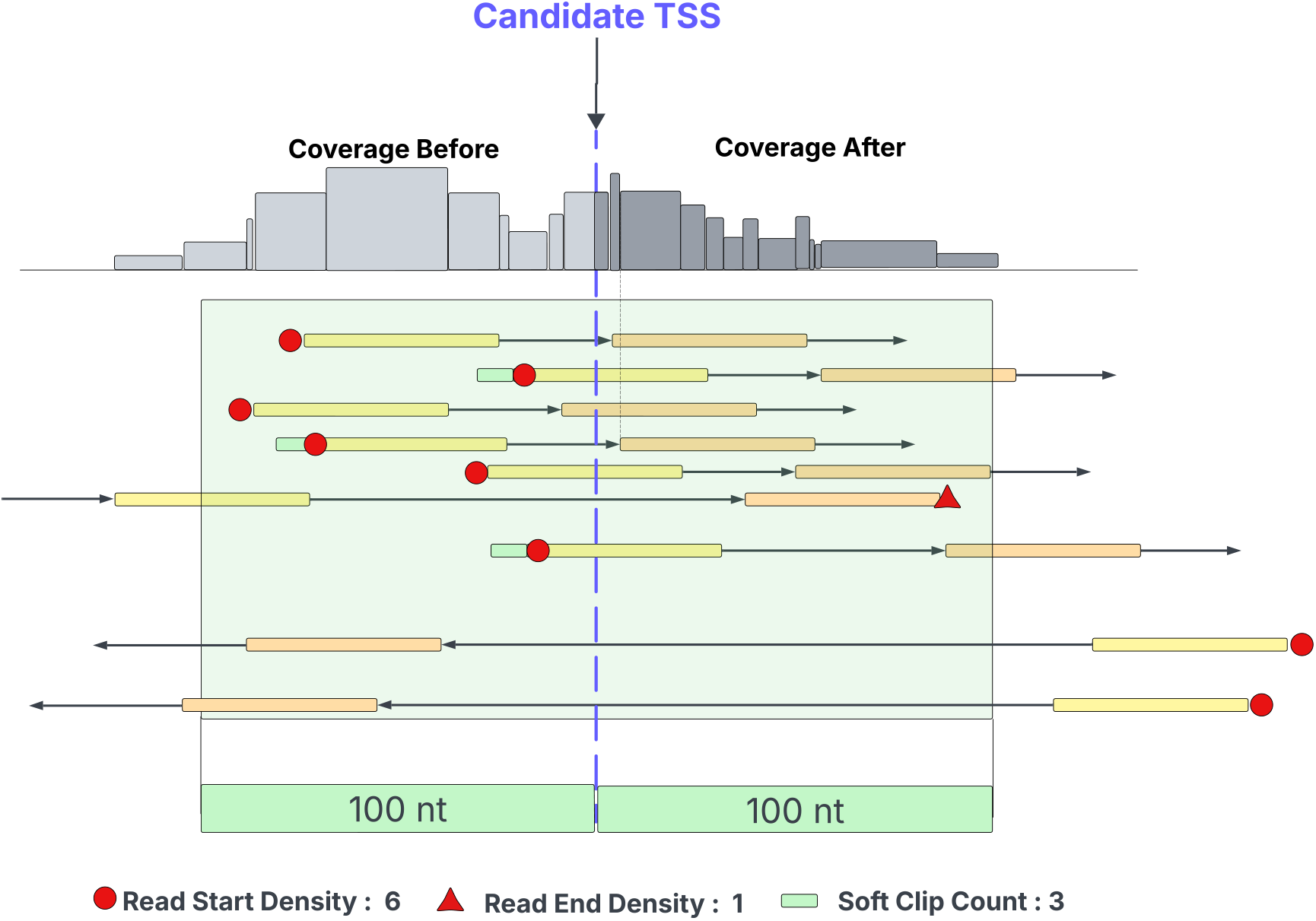
An example of feature extration. The dotted blue line marks a putative TSS. A context window of 100nt on both sides is used to extract features from the neighborhood. The top panel shows the coverage distribution. The bottom panel shows alignments of multi-exon reads, where the first and second exons are marked in yellow and orange. (Soft)-clips are marked in green. There are six reads that start, one that ends, and three reads with a soft clip in the context window.

#### Label Assignment

To enable supervised learning, each candidate TSS and TES site is assigned a binary label by comparing it to reference annotations. A candidate is labeled as true if it lies within ±50 nucleotides of an annotated TSS or TES (depending on site type), and false otherwise. This tolerance accounts for minor alignment and annotation discrepancies while preserving biological resolution. For long-reads datasets (ONT cDNA, ONT direct RNA, and PacBio), we experiment with GENCODE annotations aligned to the GRCh38 reference genome. For short-reads data, we use Ensembl gene annotations for labeling, also aligned to GRCh38. Labels are computed independently for TSS and TES datasets, allowing for site-type-specific classifiers.

#### Model Training and Evaluation

We independently train models for TSS and TES classification using two ensemble-based classifiers: XGBoost and Random Forest. For training, we use data from chromosomes 1 to 10, while data from the remaining chromosomes is reserved for testing. Once trained, each model assigns a probability score to every candidate site, representing the confidence that the site corresponds to a true TSS or TES. These site-level scores can be used independently to directly rank candidate sites, filter transcript ends, or otherwise support downstream transcript-level modeling in the second stage of Telos.

### 5.3 Stage II: Transcript-Level Scoring

While Stage I of Telos focuses on assessing individual transcript start and end sites, the second stage integrates this site-level information to evaluate the overall reliability of full-length transcript models. The goal is to distinguish plausible transcripts from likely artifacts based on both site-level confidence and features of the transcript.

#### Feature Construction

For each transcript in the input assembly, Telos constructs a feature vector by aggregating (i) TSS related features and (ii) TES related features (extracted in Stage I), (iii) TSS and TES probabilities from Stage I classification model (i.e., the classifier scores for the transcript’s 5^*′*^ and 3^*′*^ ends), (iv) abundance estimates provided by the assembler, such as TPM or coverage, and (v) basic statistics about exon lengths from the assembled transcript . They are described in Appendix B. The intuition behind including Stage I classifier scores is to explicitly incorporate site-level confidence into our assessment of full-length transcript correctness.

#### Label Assignment

To train the transcript-level model, we again use genome annotations as the source of ground truth. A transcript is labeled as true if it exactly matches an annotated isoform (class code “=” in gffcompare [18]). Transcripts with no corresponding annotation or that fall into ambiguous or unsupported categories are labeled as false. Labels are derived separately for long-read and short-read assemblies using the same annotation sources as in Stage I.

#### Models for Scoring Transcripts

Telos uses LightGBM model for transcript-level scoring. The model is trained on labeled transcripts using the features described in Section 5.3. The same train-test split procedure was used as Stage I. Once trained, the model assigns a probability to each transcript, reflecting the likelihood that it represents a full-length, biologically valid isoform. These probability scores can be used to rank assembled transcripts, prioritize high-confidence isoforms, or filter out likely artifacts.

## Funding

This work is supported by the US National Science Foundation (2145171 to M.S.) and by the US National Institutes of Health (R01HG011065 to M.S.).

## Data Availability

We used the accession numbers to identify all the datasets except cDNA NA12878 and dRNA NA12878. Both are available at https://github.com/nanopore-wgs-consortium/NA12878/blob/master/RNA.md. The short read data sets are obtained from Sequence Read Archive (SRA) with accession id: SRR307903, SRR307911, SRR315323, SRR315323, SRR315334, SRR387661, SRR534291, SRR534307, SRR534319, SRR545695, and SRR545723. Long reads data are obtained from three different projects but all of them (except the two mentioned above) are available at the ENCODE portal (https://www.encodeproject.org). PacBio datasets can be accessed using accession id: ENCFF370NFS, ENCFF450VAU, ENCFF563QZR, and ENCFF694DIE. For direct RNA datasets the accession ids are: ENCFF155CFF, ENCFF771DIX, and ERR6053059. For cDNA: ENCFF023EXJ, ENCFF263YFG, and ERR6053079.

## Code Availability

Telos is available for installation athttps://github.com/Shao-Group/Telos. All the code and script to reproduce the presented results are available at : https://github.com/Shao-Group/Telos-test.

## APPENDIX A Features in Telos Stage I

**Table 2.**
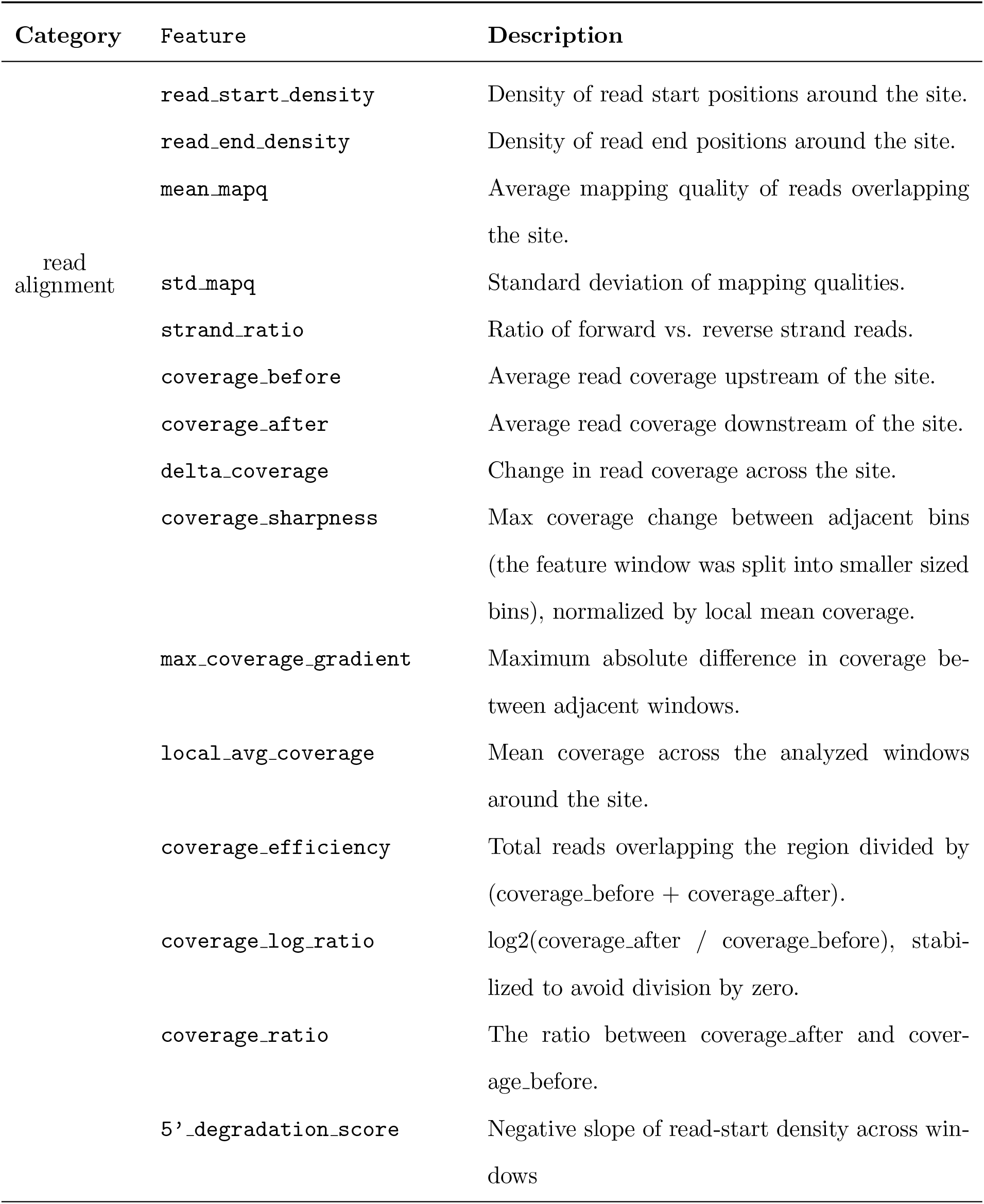
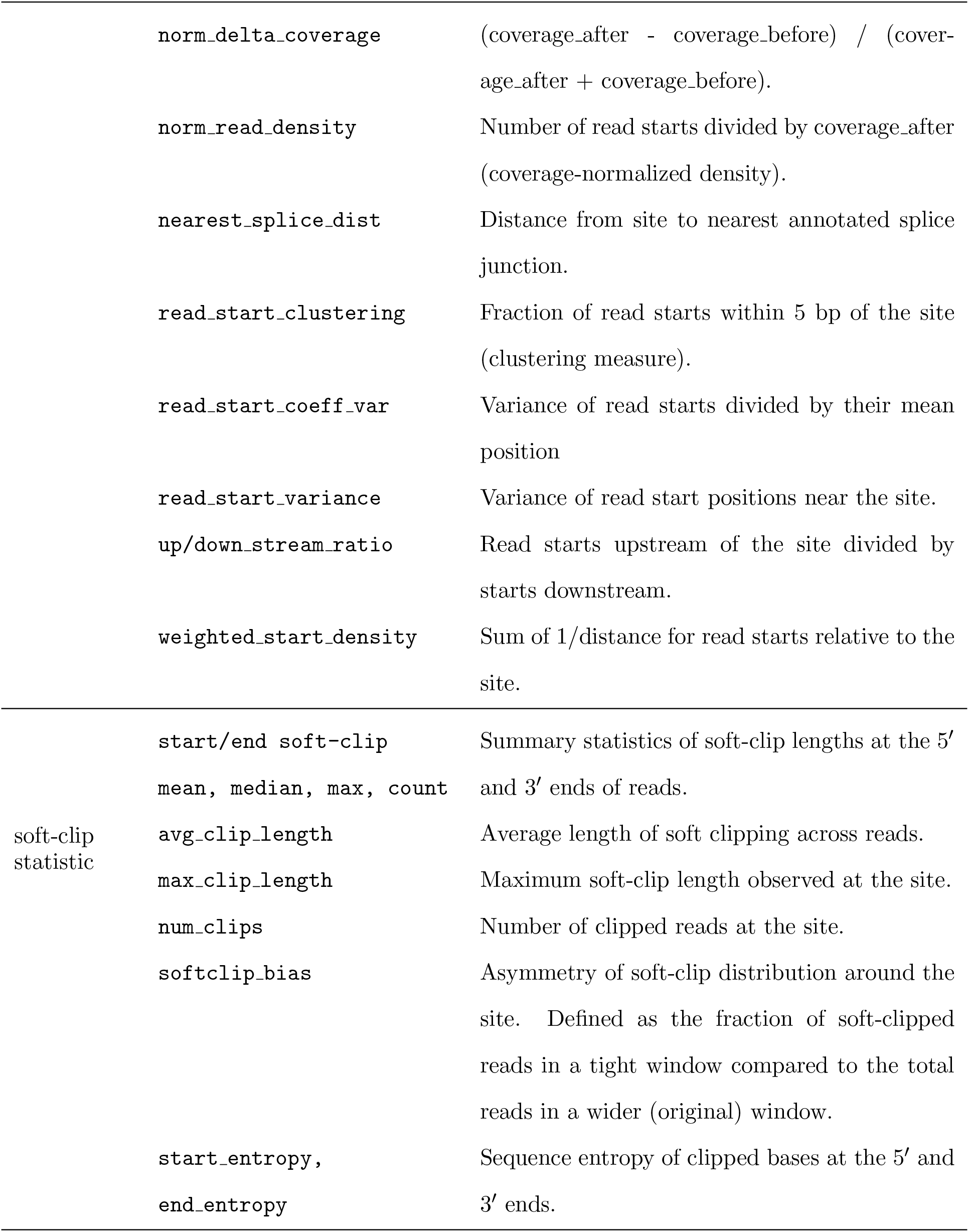
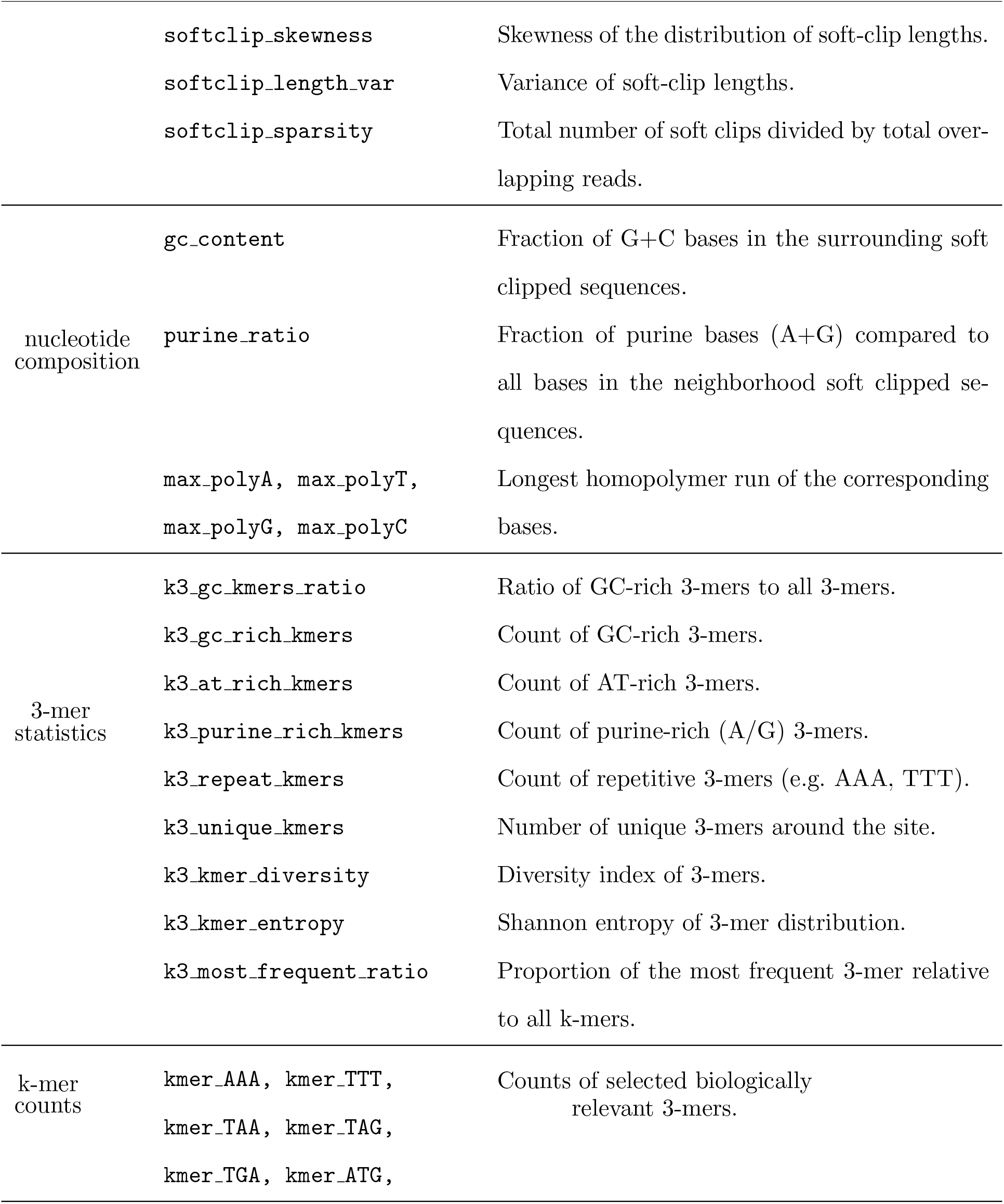
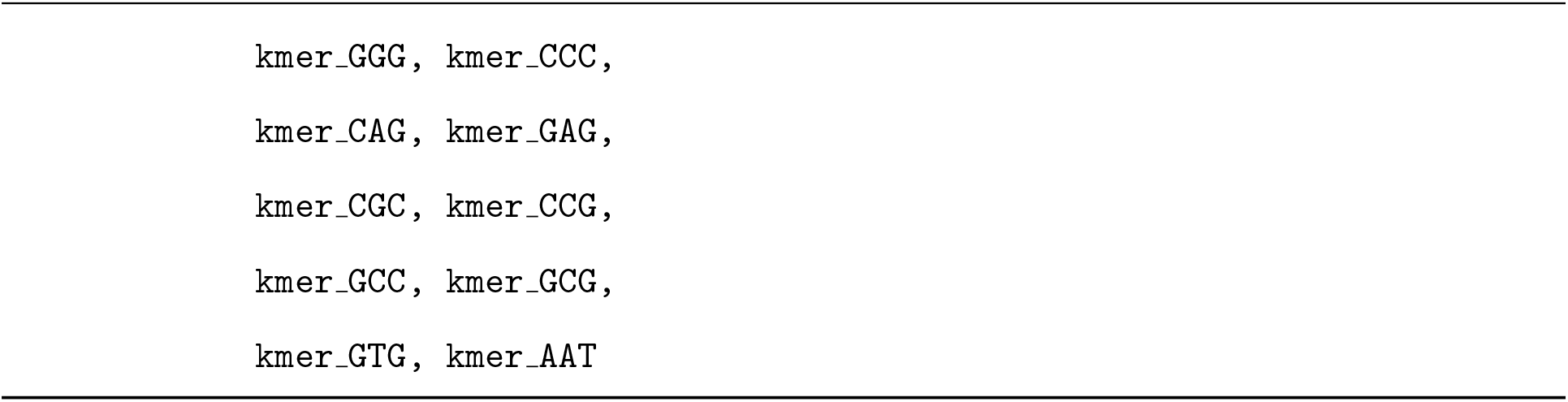
Summary of features for TSS/TES classification.

## APPENDIX B Features in Telos Stage II

**Table 3.**
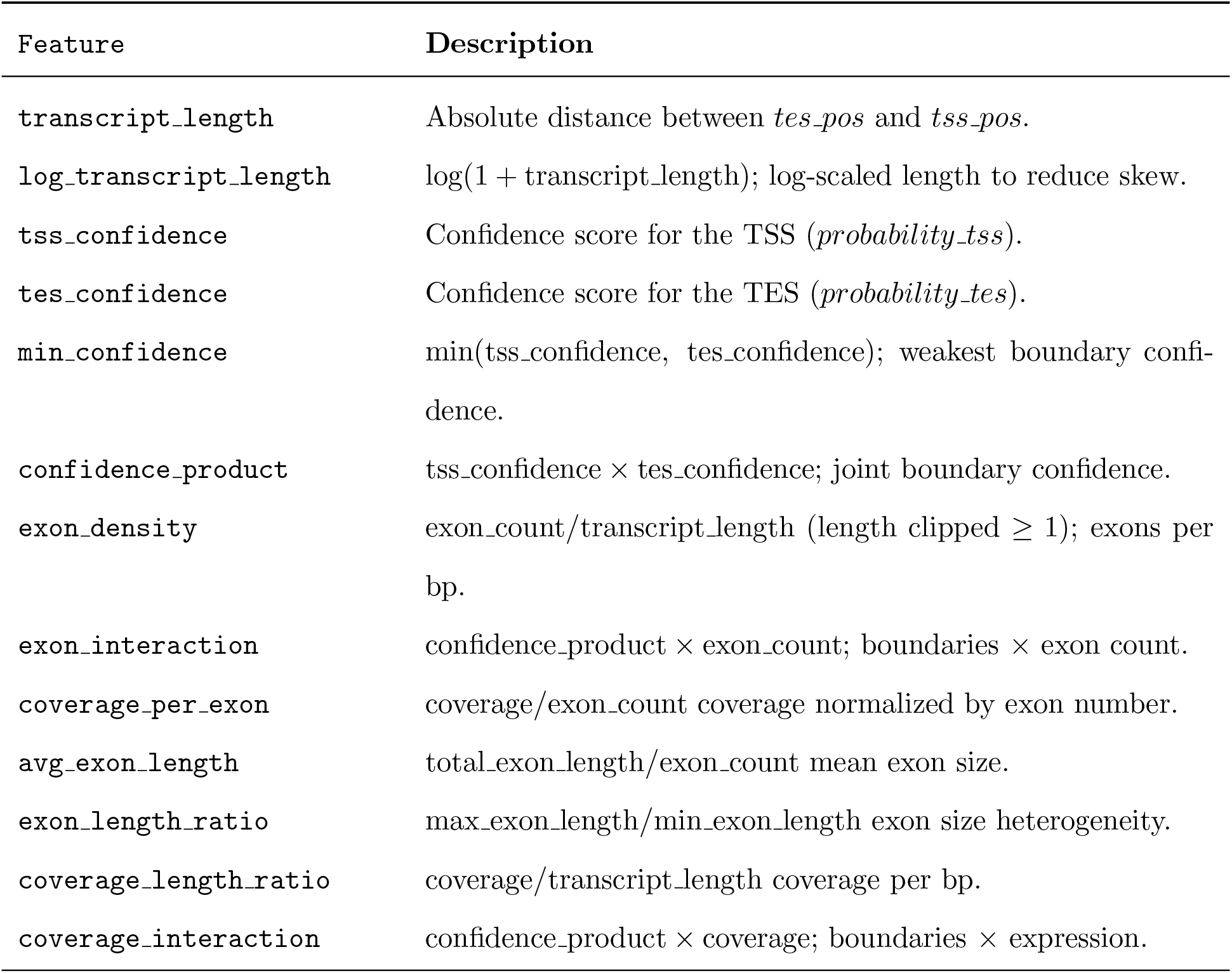
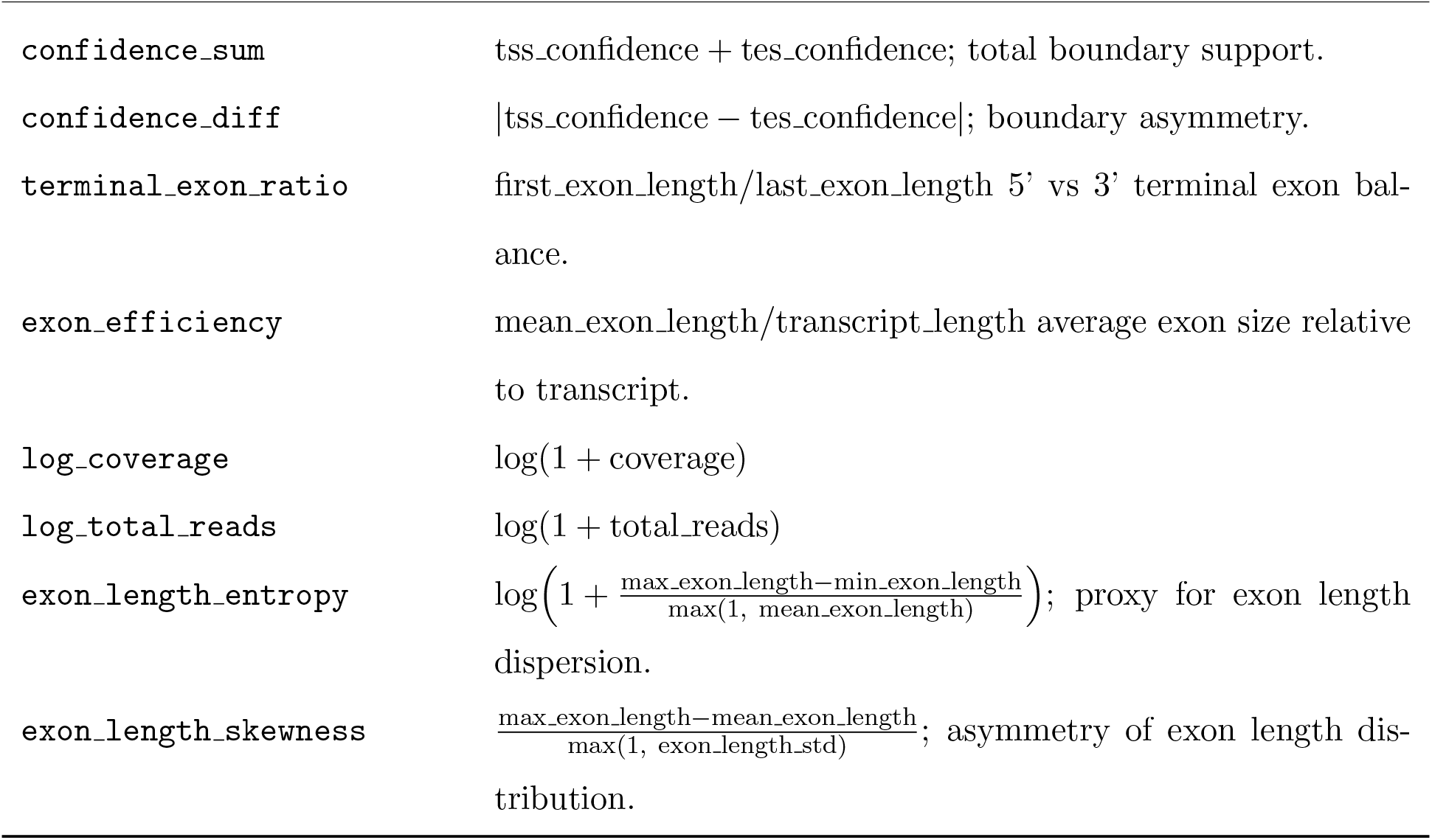
Transcript-level feature descriptions.

## Notes

### Competing Interest Statement

The authors have declared no competing interest.

